# Early life stress reduces prefrontal dendritic spine density in adult male mice

**DOI:** 10.64898/2026.11.18.767676

**Authors:** Prerna Bansal, Ernest Cote, Juan Marcos Alarcon

## Abstract

Early life stress (ELS) is a major risk factor for anxiety, depression and substance use disorders later in life, and it is thought to act in part by changing how emotion-regulating circuits develop. The prelimbic (PL) and infralimbic (IL) cortices exert top-down control over the basolateral amygdala (BLA), and this circuit is still maturing during early postnatal life. Stress in adulthood removes dendritic spines from prefrontal neurons, but it is less clear whether ELS causes lasting spine changes across this circuit, and the IL in particular has rarely been studied. Here, we asked whether ELS alters dendritic spine density in the PL, IL and BLA of adult male mice. From postnatal day (PND) 10 to 17, ELS pups were separated from the dam for 3 h per day and individually isolated from their littermates, and their home cage had reduced nesting material until PND21. Control pups were separated together with their littermates for only 15 min per day and kept standard nesting. In adulthood, brains were processed for Golgi-Cox staining, and spines were counted on proximal apical and basal dendrites of layer II/III pyramidal neurons in the PL and IL, and on dendrites of principal neurons in the BLA. ELS reduced spine density by approximately 30% in the IL and PL dendrites, with the largest losses concentrated on proximal apical dendrites. The BLA showed a smaller reduction of about 12%. These preliminary findings suggest that ELS leads to an important and lasting loss of dendritic spines in prefrontal regions that regulate emotional responses.

## 1 INTRODUCTION

Early life stress (ELS), particularly childhood maltreatment, is the leading preventable risk factor for later mental illness and substance abuse (Teicher et al., 2016). In the study of Adverse Childhood Experiences (ACE), adults who reported four or more types of childhood abuse or household dysfunction were 4 to 12 times more likely to have depression, alcoholism, drug abuse or suicide attempts, and this risk rose with each additional type of adversity (Felitti et al., 1998). These effects likely persist because ELS occurs while the brain is still developing: in rodents, the effects of stress in adulthood, even chronic stress, reverse within a few weeks, whereas the effects of severe stress early in life are long-lasting (Lupien et al., 2009). Because human studies are mostly cross-sectional and rely on adults recalling their childhood, rodent models, in which the type and timing of early stress can be controlled, are needed to link early experience to lasting brain changes (Teicher et al., 2016). In male mice, maternal separation from postnatal day 10 to 17, but not earlier in life, makes animals more susceptible to depression-like behavior after stress in adulthood, even though they show no obvious behavioral change beforehand (Peña et al., 2017).

Many of the disorders that follow ELS share a common thread: difficulty regulating fear and other emotional responses. In rodents, this regulation depends largely on the medial prefrontal cortex (mPFC) and its interactions with the basolateral amygdala (BLA) (Quirk and Mueller, 2008). In the classic model, the prelimbic cortex (PL) promotes the expression of fear, while the infralimbic cortex (IL) helps consolidate and retrieve extinction memories and can dampen amygdala output (Quirk and Mueller, 2008). Inactivating PL reduces fear expression without affecting extinction memory, inactivating IL impairs extinction without affecting fear expression, and inactivating the BLA impairs both (Sierra-Mercado et al., 2011); silencing IL inputs to the amygdala also impairs extinction (Bukalo et al., 2015; Bloodgood et al., 2018). In mice, PL forms strong reciprocal connections with the anterior BLA, whereas IL receives its BLA input mainly from the caudal BLA (Hintiryan et al., 2021; Manoocheri and Carter, 2022). This circuit also matures late. In the mouse mPFC, excitatory inputs onto layer 3 pyramidal neurons increase rapidly during the second postnatal week (Kroon et al., 2019; Chen et al., 2025), and connections between the mPFC and BLA continue to develop into adolescence, maturing differently for PL and IL (Pattwell et al., 2016; Arruda-Carvalho et al., 2017). A circuit still under construction this early in life may be especially sensitive to adversity: in children who experienced early maternal deprivation, amygdala-mPFC connectivity takes on an adult-like pattern ahead of schedule (Gee et al., 2013).

Dendritic spines offer a direct way to see how early experience reshapes this circuit. Each spine typically receives a single excitatory synapse (Nimchinsky et al., 2002), so spine density gives an estimate of how much excitatory input a neuron can receive, and because different parts of the dendritic tree receive distinct inputs, where spines are lost can matter as much as how many (Spruston, 2008). Spines are also directly sensitive to stress hormones: in living mice, corticosterone rapidly increases spine turnover, and chronic glucocorticoid excess removes stable spines formed early in life (Liston and Gan, 2011). In adult rats, three weeks of daily restraint stress shortens apical, but not basal, dendrites and lowers apical spine density by 16% in layer II/III pyramidal neurons of the anterior cingulate and prelimbic cortices (Radley et al., 2004; Radley et al., 2006), whereas in the BLA chronic immobilization stress enlarges dendritic arbors (Vyas et al., 2002). ELS seems to push the mPFC in the same direction as adult stress: maternal separation, including separation with individual isolation, and limited bedding all reduce dendritic length or spine density in the prelimbic and medial prefrontal cortex, in some cases only after puberty (Monroy et al., 2010; Chocyk et al., 2013; Xu et al., 2023). These changes matter for behavior: in mice, loss of PL spines after early adversity impairs spatial working memory (Xu et al., 2023), and in rats, IL dendritic and spine features after acute stress predict how well fear extinction is later retrieved (Moench et al., 2016). Still, the direction of these effects depends on when during development the stress occurs, and maternal separation has also been reported to increase spine density in the anterior cingulate cortex (Bock et al., 2005; Gos et al., 2008; Muhammad and Kolb, 2011). The BLA is even less settled: maternal separation in rats increases BLA dendritic size and spine density in adulthood (Koe et al., 2016), whereas restraint stress in juvenile rats reduces BLA spines (Pinzón-Parra et al., 2019). The IL has received the least attention; one of the few studies found that maternal separation lengthened IL apical dendrites and reduced thin spines in adolescent females but not males (Farrell et al., 2016). Few studies have examined IL spines in adulthood after ELS, and no study has yet compared PL, IL and BLA within the same animals.

Here, we asked whether ELS alters dendritic spine density in the PL, IL and BLA of adult male mice. Given the role of the PL and IL in top-down control of emotional responses, and that ELS reduces prefrontal spine density, we predicted that ELS would lower spine density more strongly in the mPFC than in the BLA. Mice were exposed to maternal separation with individual isolation and reduced nesting material from postnatal day 10 to 17. In adulthood, we used Golgi-Cox staining to quantify spines on proximal apical and basal dendrites of layer II/III pyramidal neurons in the PL and IL, and on the dendrites of principal neurons in the BLA, sampling all three regions within the same animals. We found that ELS reduced spine density, with the largest reductions in the IL and PL, particularly on proximal apical dendrites.

## 2 METHODS

### 2.1 Animals

C57BL/6 wild-type breeding pairs were maintained under standard laboratory conditions to generate the litters used in this study. Animals were housed in a temperature- and humidity-controlled facility on a 12-hour light/dark cycle with food and water available ad libitum. Litters were assigned to either the ELS or the control condition. All pups in a litter, male and female, underwent the procedure assigned to that litter. Pups were weaned at postnatal day (PND) 21 and then group-housed with same-sex cagemates under identical conditions, undisturbed except for routine cage changes, until adulthood. Only male offspring were used. Animals came from two cohorts and four litters, with two litters per group. All procedures were approved by the SUNY Downstate Health Sciences University Institutional Animal Care and Use Committee (IACUC protocol 20-10578).

### 2.2 Early life stress protocol

ELS was induced from PND10 to PND17 using a protocol adapted from previous work in our laboratory (Cote, 2023; Fig. 1A). This late postnatal window was chosen because stress during it, unlike stress earlier in life, increases susceptibility to stress in adulthood in male mice (Peña et al., 2017). Each day, in the morning during the light phase, pups in the ELS group were separated from the dam for 3 hours and individually isolated, with each pup placed in its own enclosure under a dome so that it had no physical contact with either the dam or its littermates. To maintain body temperature, pups were kept on a heating surface during separation, their temperature was monitored so that it did not fall below 27 °C, and they were checked periodically. In addition, half of the nesting material was removed from ELS home cages on PND10 and withheld until PND21, to increase maternal stress and discourage the dam from compensating with extra care (Cote, 2023). The control group consisted of pups separated for 15 minutes per day from PND10 to PND17, kept together with their littermates to allow normal huddling while controlling for maternal absence, and housed with standard nesting material. After each session, pups were returned to the home cage. Because brief daily separation resembles early handling, which by itself can change later stress responses (Plotsky and Meaney, 1993), control pups were not an undisturbed group. ELS and control pups therefore differed in three ways: the length of daily separation (3 h vs. 15 min), the loss of contact with littermates during separation, and reduced nesting material from PND10 to PND21.

**Figure 1.**
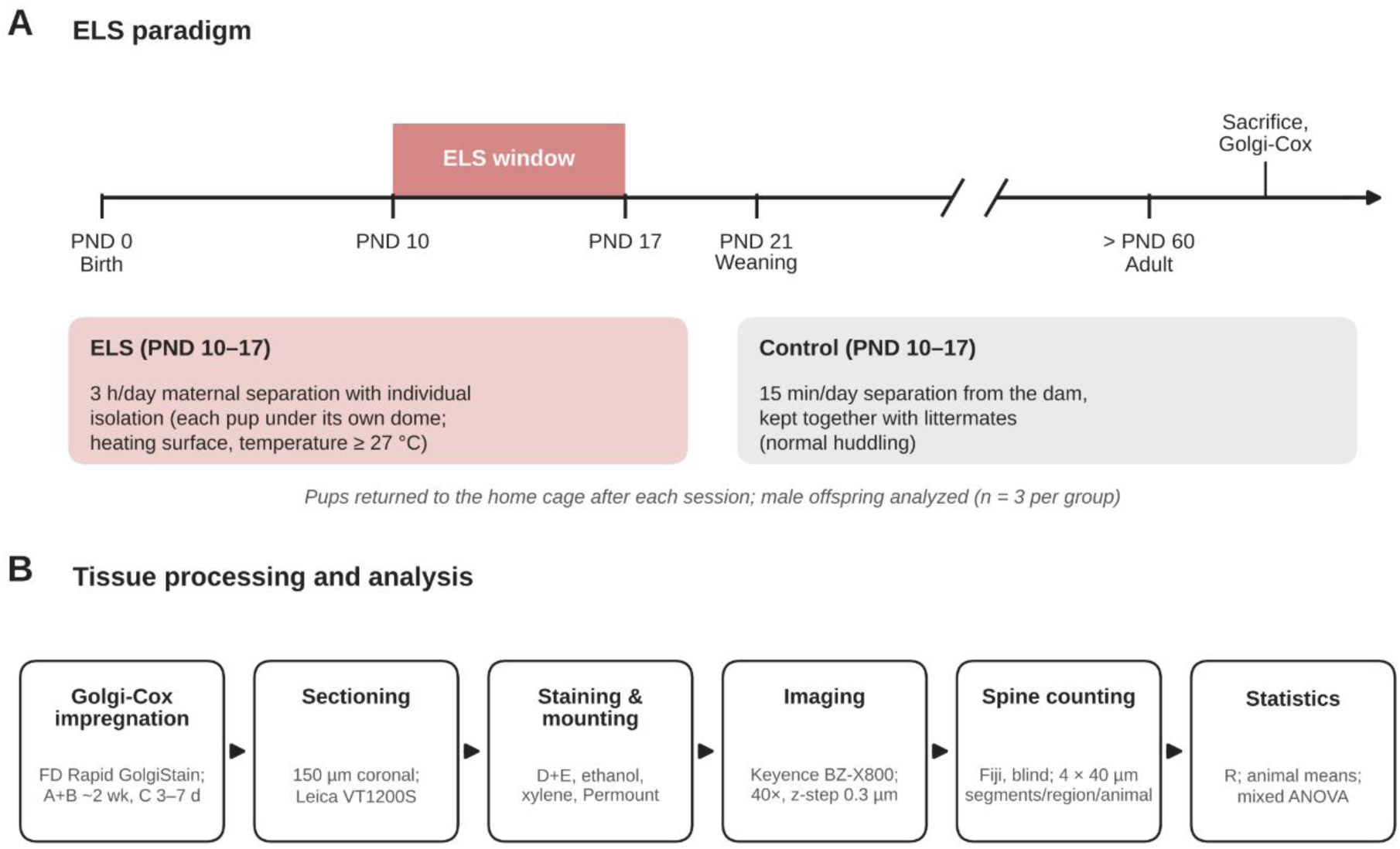
Experimental design. **(A)** Timeline of the early life stress (ELS) paradigm. From postnatal day (PND) 10 to PND 17, ELS pups were separated from the dam for 3 h per day and individually isolated, and their home cage had reduced nesting material until PND 21, whereas control pups were separated for 15 min per day and kept together with their littermates. Pups were returned to the home cage after each session and weaned at PND 21. Male offspring were sacrificed in adulthood, after PND 60, for Golgi-Cox staining (n = 3 per group). **(B)** Tissue processing and analysis: Golgi-Cox impregnation, 150 µm coronal sectioning, staining and mounting, imaging at 40× with a 0.3 µm z-step, blinded spine counting in Fiji, and statistical analysis of animal means in R.

### 2.3 Golgi-Cox staining and tissue preparation

Mice were sacrificed as adults, after PND 60 (Fig. 1A). They were deeply anesthetized with isoflurane (5% vaporized in 100% oxygen) until loss of the righting reflex and then euthanized by rapid decapitation. Brains were rapidly removed and one hemisphere from each brain was processed for Golgi-Cox staining; left and right hemispheres were both represented across animals to avoid a hemisphere bias. Tissue was processed with the FD Rapid GolgiStain Kit (FD NeuroTechnologies, Columbia, MD), following the staining manual provided with the kit and established laboratory protocols (Fig. 1B). Tissue was immersed in the impregnation solutions A and B (FD Rapid GolgiStain Kit) and kept in the dark at room temperature for approximately two weeks, and then transferred to Solution C (FD Rapid GolgiStain Kit), where it was kept in the dark for 3 to 7 days before sectioning. Coronal sections were cut at 150 µm on a vibratome (Leica VT1200S) and mounted on gelatin-coated slides. Mounted sections were rinsed in distilled water, developed in a mixture of Solutions D and E (FD Rapid GolgiStain Kit), dehydrated through a graded ethanol series, cleared in xylene and coverslipped with Permount mounting medium and glass coverslips. Slides were left to dry for two weeks before imaging.

### 2.4 Regions of interest

Three regions were analyzed in each animal: the prelimbic (PL) and infralimbic (IL) cortices of the mPFC and the basolateral amygdala (BLA). Regional boundaries were identified using the coronal Allen Mouse Brain Reference Atlas (Allen Institute for Brain Science, 2011), with sections matched to atlas images 27–39 for the PL, 34–45 for the IL and 63–75 for the BLA (Fig. 2A). In the PL and IL, analysis was restricted to pyramidal neurons in layers II/III. Layers II/III were chosen because excitatory inputs onto layer 3 pyramidal neurons of the mouse mPFC increase rapidly during the second postnatal week, overlapping the ELS window (Kroon et al., 2019; Chen et al., 2025), and because stress-induced dendritic and spine loss in the mPFC has been described in these layers (Radley et al., 2004; Radley et al., 2006). These layers were identified as the dense band of pyramidal cell bodies just below the cell-sparse layer I (Leuner and Gould, 2010), and neurons were included only if their soma lay within these layers and within the boundaries of the PL or IL (Fig. 2B, C). In the BLA, pyramidal-like principal neurons were selected; because these neurons have a consistent apical dendrite only in the caudal part of the posterior basolateral nucleus (McDonald, 1984), BLA dendrites were not classified as apical or basal and were pooled for analysis.

**Figure 2.**
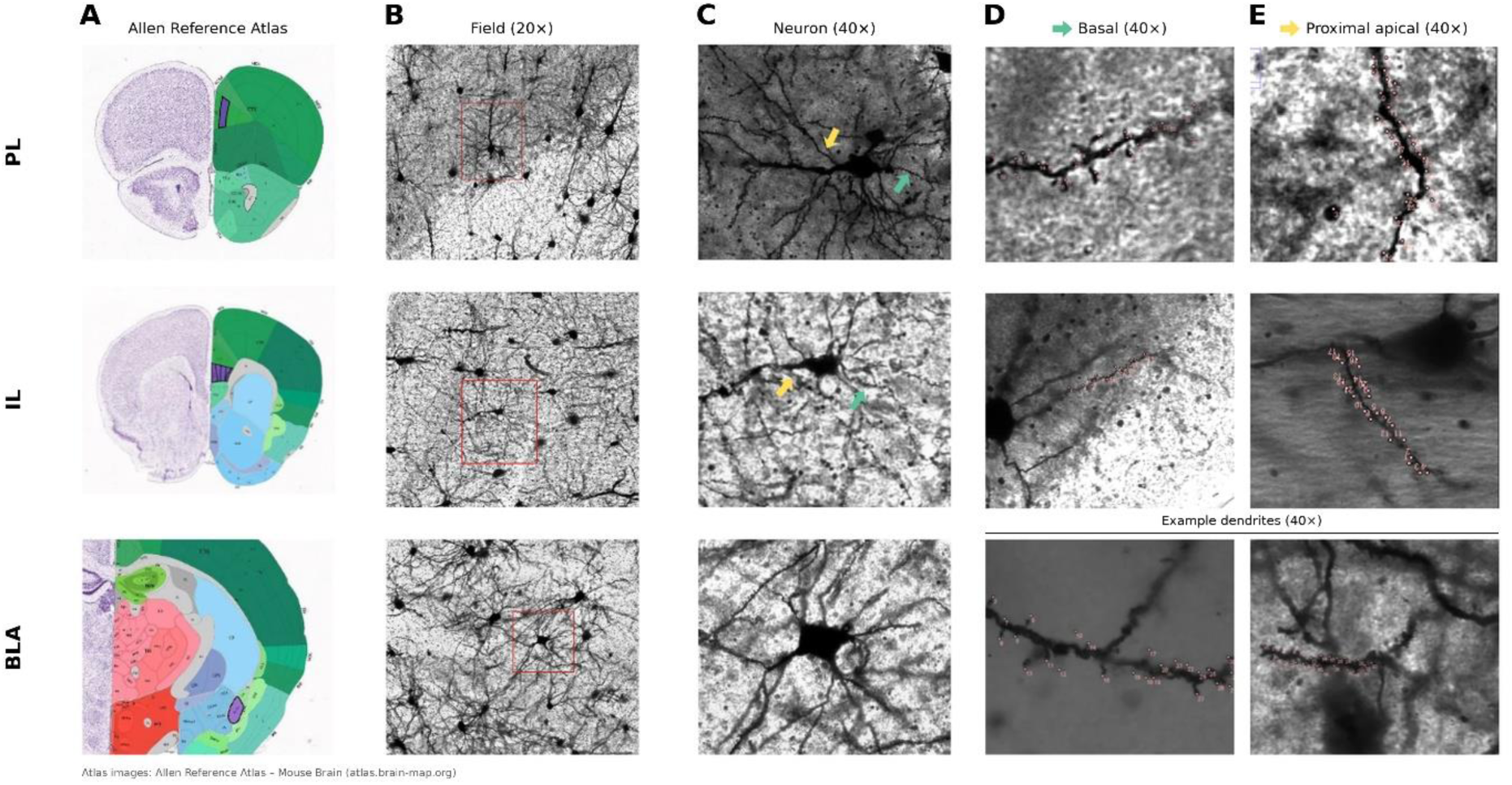
Regions, neurons and dendrites sampled for spine analysis. **(A)** Coronal sections from the Allen Reference Atlas showing the location of the PL, IL and BLA (purple; Allen Institute for Brain Science, 2011). **(B)** Golgi-Cox-stained fields (20×) with an example neuron boxed in red. **(C)** Example neurons (40×); green arrow, basal dendrite; yellow arrow, proximal apical dendrite. **(D, E)** Example counted segments (40×) from basal **(D)** and proximal apical (E) dendrites of PL and IL pyramidal neurons. BLA dendrites, which were not divided by dendrite type, are shown as example dendrites of principal neurons. Dots mark counted spines. Images are examples for illustration and are not intended to represent group differences.

### 2.5 Imaging

Golgi-stained sections were imaged under brightfield illumination on a Keyence BZ-X800 all-in-one microscope (Keyence, Osaka, Japan). Z-stacks of selected dendritic segments were acquired with a 40× Plan Apochromat objective (NA 0.95, dry) using BZ-X800 Viewer software, with a z-step of 0.3 µm so that spines could be followed across focal planes. Imaging settings were kept constant across all animals. Similar 40× imaging has been used to quantify spine density on Golgi-stained dendrites, including apical oblique dendrites of pyramidal neurons in the PL and IL (Jones et al., 2021) and dendrites in the mouse frontal cortex (Weyer et al., 2026). Only visible spines were counted, and no correction was made for spines hidden behind the dendritic shaft, an approach commonly used when comparing groups (Bock et al., 2005).

### 2.6 Neuron and dendrite selection

In the PL and IL, pyramidal neurons were identified by their triangular soma, a single prominent apical dendrite and a basal dendritic arbor; BLA principal neurons were identified by their pyramidal-like soma and spiny dendrites. Neurons were included only if they were fully impregnated, free of staining artifacts and clearly separated from neighboring cells. In each region of each animal, four 40 µm dendritic segments were analyzed (Fig2D,E), each taken from a different neuron, and every neuron came from a different section. In total, 12 neurons were analyzed per animal (72 across the study), and in the PL and IL each dendrite-type mean was based on two segments per animal. In the PL and IL, these were two apical and two basal segments. Apical segments were taken from oblique branches arising from the apical trunk close to the soma, often called apical oblique dendrites (Li et al., 2023), and are referred to here as proximal apical dendrites. Basal segments were taken from primary basal dendrites arising directly from the soma. In the BLA, four segments were sampled without regard to dendrite type. Because spine density varies with position on the dendritic tree, segments were taken from consistent locations within each dendrite type (Li et al., 2023).

This is particularly important for apical dendrites of mPFC pyramidal neurons, where spines are sparse on the proximal trunk and spine density changes with distance from the soma (Radley et al., 2006). Like Radley et al. (2006), who sampled small second- and third-order branches arising from the apical trunk near the soma, we took apical segments from this same type of branch in every neuron, keeping this source of variation similar across neurons and groups. Segments were chosen to avoid branch points, truncated endings and overlapping dendrites.

### 2.7 Spine quantification

Z-stacks were opened in Fiji (ImageJ; Schindelin et al., 2012), the spatial scale was set from the image calibration, and spines were counted manually by scrolling through the full stack, using the same display settings for all images, with the counter blind to group. A protrusion was counted as a spine if it extended clearly from the dendritic shaft and was visible across adjacent focal planes; ambiguous protrusions and staining artifacts were excluded. Spines were counted along each 40 µm segment, and spine density was expressed as spines per 10 µm of dendrite to allow comparison with previous studies. Segment values were averaged within each region (and, in the PL and IL, within each dendrite type) for each animal, so that the animal, rather than the neuron or segment, was the unit of analysis, avoiding pseudoreplication (Li et al., 2023). To keep the analysis blind, each mouse was assigned a lab ID after weaning that did not indicate its group; neurons were selected, imaged and counted using these IDs, which were matched to group only after all counts were complete.

### 2.8 Statistical analysis

All analyses were run in R (version 4.3.2) using the afex package (version 1.3-1) for ANOVA and the effectsize package (version 0.8.7) for effect sizes (R Core Team, 2023; Singmann et al., 2024; Ben-Shachar et al., 2020). Two mixed-design ANOVAs were performed on animal means. The first tested Group (control, ELS) as a between-subjects factor and Region (PL, IL, BLA) as a within-subjects factor, using spine density pooled across dendrites. The second, restricted to the prefrontal regions, tested Group as a between-subjects factor and Region (PL, IL) and Dendrite (proximal apical, basal) as within-subjects factors. Greenhouse–Geisser corrections were applied to within-subjects effects with more than two levels (Region in the first ANOVA). To describe the pattern of effects, ELS and control animals were then compared overall, within each region and, in the PL and IL, within each dendrite type, using Welch’s t-tests. Because these follow-up comparisons were exploratory, their p values are reported without correction for multiple comparisons. For the ANOVAs, effect sizes are reported as generalized eta squared (η^2^G; Olejnik and Algina, 2003; Bakeman, 2005), as computed by afex; for the follow-up comparisons, they are reported as percent change from control and Cohen’s d (calculated with the pooled standard deviation). Statistical significance was set at p < .05, and data are presented as mean ± SEM. With three animals per group, normality could not be meaningfully tested. Figures were prepared in R with the ggplot2 package (Wickham, 2016).

## 3 RESULTS

### 3.1 Overall spine density across regions

To test whether ELS alters spine density in adulthood, we first compared the ELS and control groups across the PL, IL and BLA (n = 3 mice per group). A mixed ANOVA revealed a main effect of group, F(1, 4) = 19.24, p = .012, η^2^G = .596: ELS mice had fewer spines than controls. Averaged across all three regions, spine density was 23.7% lower in ELS mice than in controls (5.55 vs. 7.27 spines/10 µm; Welch’s t-test, t(2.59) = −4.39, p = .029, d = −3.58; Fig. 3; Table 1). Neither the effect of region, F(1.57, 6.28) = 0.44, p = .616, η^2^G = .071, nor the group × region interaction, F(1.57, 6.28) = 1.14, p = .361, η^2^G = .165, was significant (Greenhouse–Geisser corrected), so at this sample size the size of the ELS effect could not be shown to differ between regions.

**Figure 3.**
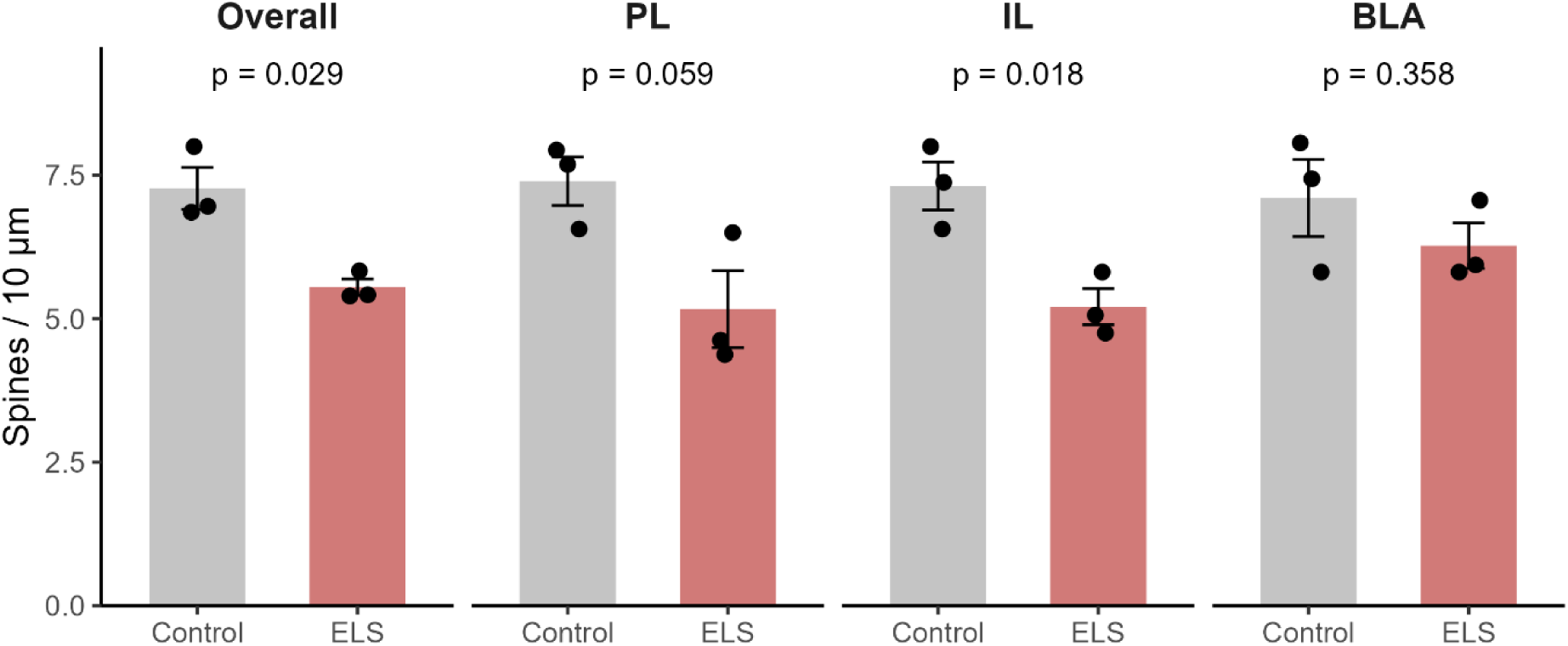
ELS reduces dendritic spine density, with the largest reductions in the prefrontal cortex. Spine density (spines per 10 µm) averaged across all regions (Overall) and in the PL, IL and BLA of control and ELS mice. BLA dendrites were pooled across dendrite type. Bars show mean ± SEM; dots show individual animals (n = 3 per group). p values are from Welch’s t-tests; comparisons for individual regions are exploratory and uncorrected for multiple comparisons.

**Table 1.**
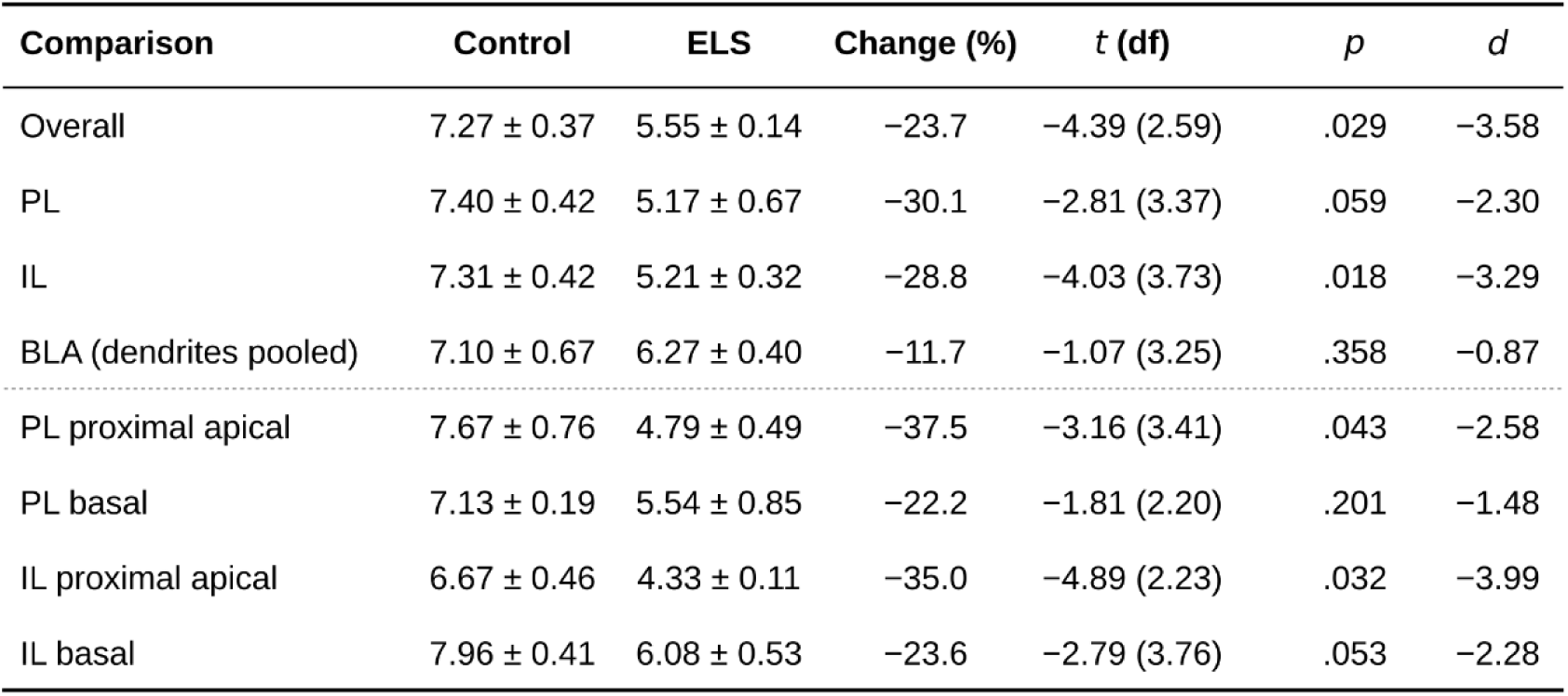
Spine density in control and ELS mice. Values are mean ± SEM spines per 10 µm (n = 3 animals per group). t values, with Welch-corrected degrees of freedom in parentheses, and p values are from Welch’s t-tests; comparisons below the overall row are exploratory and uncorrected. d, Cohen’s d.

| **Comparison** | **Control** | **ELS** | **Change (%)** | ***t* (df)** | *p* | *d* |
|---|---|---|---|---|---|---|
| Overall | 7.27 ± 0.37 | 5.55 ± 0.14 | −23.7 | −4.39 (2.59) | .029 | −3.58 |
| PL | 7.40 ± 0.42 | 5.17 ± 0.67 | −30.1 | −2.81 (3.37) | .059 | −2.30 |
| IL | 7.31 ± 0.42 | 5.21 ± 0.32 | −28.8 | −4.03 (3.73) | .018 | −3.29 |
| BLA (dendrites pooled) | 7.10 ± 0.67 | 6.27 ± 0.40 | −11.7 | −1.07 (3.25) | .358 | −0.87 |
| PL proximal apical | 7.67 ± 0.76 | 4.79 ± 0.49 | −37.5 | −3.16 (3.41) | .043 | −2.58 |
| PL basal | 7.13 ± 0.19 | 5.54 ± 0.85 | −22.2 | −1.81 (2.20) | .201 | −1.48 |
| IL proximal apical | 6.67 ± 0.46 | 4.33 ± 0.11 | −35.0 | −4.89 (2.23) | .032 | −3.99 |
| IL basal | 7.96 ± 0.41 | 6.08 ± 0.53 | −23.6 | −2.79 (3.76) | .053 | −2.28 |

### 3.2 Spine density by region

Exploratory Welch’s t-tests within each region, uncorrected for multiple comparisons, showed that the largest reductions were in the prefrontal cortex. Spine density was 28.8% lower in the IL of the ELS group (5.21 vs. 7.31 spines/10 µm; t(3.73) = −4.03, p = .018, d = −3.29) and 30.1% lower in the PL, although this difference was not statistically significant (5.17 vs. 7.40; t(3.37) = −2.81, p = .059, d = −2.30). In the BLA, by contrast, ELS was associated with a smaller, nonsignificant reduction of 11.7% (6.27 vs. 7.10; t(3.25) = −1.07, p = .358, d = −0.87; Fig. 3; Table 1).

### 3.3 Spine density by region and dendrite type

We next asked whether the loss of spines in the PL and IL differed between proximal apical and basal dendrites (Fig. 4). A second mixed ANOVA restricted to these regions confirmed the main effect of group, F(1, 4) = 15.82, p = .016, η^2^G = .674, and showed that, averaged across regions, basal dendrites carried more spines than proximal apical dendrites in both groups, F(1, 4) = 14.08, p = .020, η^2^G = .225. The group × dendrite interaction was not significant, F(1, 4) = 4.08, p = .113, η^2^G = .078. The remaining effects were not significant (region × dendrite, F(1, 4) = 7.09, p = .056, η^2^G = .181; all others, p ≥ .477).

**Figure 4.**
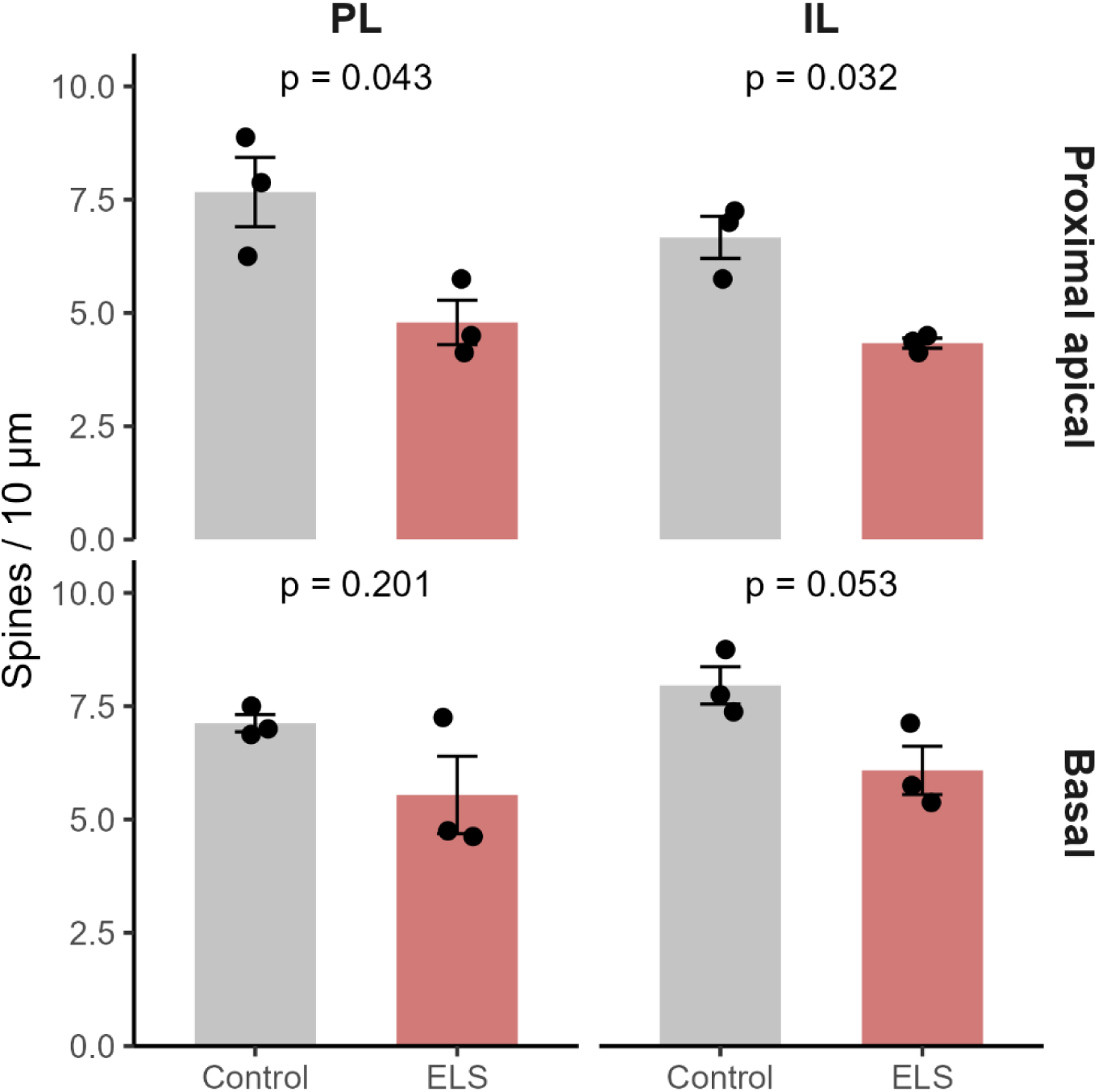
Spine loss in the PL and IL by dendrite type. Spine density (spines per 10 µm) on proximal apical and basal dendrites of layer II/III pyramidal neurons in the PL and IL of control and ELS mice. Bars show mean ± SEM; dots show individual animals (n = 3 per group). p values are from exploratory Welch’s t-tests, uncorrected for multiple comparisons.

Exploratory Welch’s t-tests nonetheless showed that the largest losses were on proximal apical dendrites, where spine density was 37.5% lower in the PL (4.79 vs. 7.67 spines/10 µm; t(3.41) = −3.16, p = .043, d = −2.58) and 35.0% lower in the IL (4.33 vs. 6.67; t(2.23) = −4.89, p = .032, d = −3.99) of ELS mice. Reductions on basal dendrites were smaller, at 23.6% in the IL (6.08 vs. 7.96; t(3.76) = −2.79, p = .053, d = −2.28) and 22.2% in the PL (5.54 vs. 7.13; t(2.20) = −1.81, p = .201, d = −1.48; Fig. 4; Table 1). Because these follow-up comparisons were not corrected for multiple comparisons, they should be read as describing the pattern of the main effect rather than as independent findings.

## 4 DISCUSSION

In this preliminary study, ELS, combining maternal separation with individual isolation and reduced nesting material from PND10 to PND17, lowered dendritic spine density in adult male mice. Across the PL, IL and BLA, ELS mice had about 24% fewer spines than controls. The reduction was largest in the prefrontal cortex, where spine density fell by 29% in the IL and 30% in the PL, with the greatest losses, 35–38%, on proximal apical dendrites. The BLA showed a smaller reduction of about 12%. Together, these data point to the IL and PL, the regions that exert top-down control over emotional responses, as the parts of this circuit most affected by early adversity. By sampling the PL, IL and BLA within the same animals, this study adds the IL, a region rarely examined after ELS, to the prefrontal areas known to be structurally altered by early adversity. Because these results come from three animals per group, we see this regional pattern as a lead to confirm in a larger sample.

### 4.1 Prefrontal spine loss after ELS in the context of previous work

The prefrontal spine loss we observed fits with earlier work on early adversity. Limited bedding and nesting from PND2 to PND9 reduces spine density on apical, but not basal, dendrites of PL pyramidal neurons in young adult mice (Xu et al., 2023), maternal separation lowers apical and basal spine density in layer II/III pyramidal neurons of the mPFC in adolescent rats (Chocyk et al., 2013), and repeated isolation from the dam and littermates reduces prefrontal spine density in rats after puberty (Monroy et al., 2010). Our results extend these findings to the IL, which has rarely been examined after ELS, and show that the loss is still present well into adulthood. Not every study agrees, however. Separation in which pups stay together with their littermates has been reported to increase spine density in the anterior cingulate cortex (Muhammad and Kolb, 2011), and in the anterior cingulate the direction of the change depends on when during development separation occurs (Bock et al., 2005; Gos et al., 2008). Differences in the region examined, the timing of the stressor, whether pups were isolated from their littermates, and the age at testing are all likely to contribute. Age may be especially relevant. Monroy et al. (2010) found that prefrontal spine loss after separation appeared only after puberty, and Farrell et al. (2016) reported IL changes after maternal separation in adolescent females but not males. The clear IL reduction we see in adult males raises the possibility that some effects of ELS on the IL take time to emerge.

### 4.2 Proximal apical versus basal dendrites

Within the PL and IL, the reductions were largest on proximal apical dendrites, where spine density fell by 35–38%, compared with 22–24% on basal dendrites. This apical emphasis is consistent with earlier work. In adult rats, chronic restraint stress removes spines and shortens dendrites on the apical, but not the basal, side of PL and anterior cingulate pyramidal neurons (Radley et al., 2004; Radley et al., 2006), and limited bedding and nesting in mice reduces spines on PL apical dendrites while sparing basal ones (Xu et al., 2023). In both studies the losses were greatest on distal apical segments, around 200 µm or more from the soma (Radley et al., 2006; Xu et al., 2023). We sampled proximal oblique branches near the soma and still found the strongest reductions on apical dendrites, which suggests that the effect of ELS on apical dendrites is not limited to their distal parts. Different long-range inputs target different parts of the dendritic tree: in the mouse mPFC, inputs from the midline thalamus, contralateral mPFC, BLA and ventral hippocampus each contact distinct populations of spines on layer 2 pyramidal neurons (Little and Carter, 2012), and BLA inputs preferentially target large spines on proximal dendrites (Little and Carter, 2013). A loss concentrated on proximal apical dendrites could therefore reflect the weakening of specific inputs, for example from the BLA, rather than a uniform reduction in excitatory input. Golgi staining cannot identify which inputs the lost spines received, so combining spine analysis with input-specific labeling would be needed to test this.

### 4.3 A smaller effect in the BLA

The BLA showed a smaller reduction of about 12%. Although not statistically significant, the direction of this change is notable because stress in the BLA is often associated with the opposite: chronic immobilization stress in adult rats enlarges BLA dendrites (Vyas et al., 2002), and maternal separation in rats increases BLA dendritic size and spine density in adulthood (Koe et al., 2016). The BLA literature is not uniform, though. The direction of BLA remodeling depends on the type of stressor (Vyas et al., 2002), and restraint stress in juvenile rats reduces BLA spine density (Pinzón-Parra et al., 2019). Our ELS protocol, which combined separation from the dam with isolation from littermates and reduced nesting material in a narrow postnatal window, may therefore engage the BLA differently from the paradigms that produce dendritic hypertrophy, that is, larger dendritic arbors and higher spine density (Vyas et al., 2002; Koe et al., 2016). Methodological factors also shape what we can say here. BLA principal neurons lack a consistent apical dendrite across most of the nucleus (McDonald, 1984), so we pooled BLA dendrites, and BLA staining was denser and harder to sample, which likely added variability in this region. A larger sample will be needed to tell whether the BLA follows the same pattern as the prefrontal cortex.

### 4.4 Possible functional consequences

What might fewer prefrontal spines mean for behavior? Because each spine typically receives a single excitatory synapse (Nimchinsky et al., 2002), a 30% loss in the IL and PL suggests that pyramidal neurons in these regions have fewer sites at which to receive excitatory input. The IL is needed to acquire and retrieve extinction memories and is thought to dampen amygdala output (Quirk and Mueller, 2008; Sierra-Mercado et al., 2011), and in adult rats exposed to a single acute stressor, IL spine density and spine type track how well animals learn and retrieve extinction (Moench et al., 2016). In the PL, spine loss after early adversity is linked to poorer working memory, and boosting PL pyramidal cell activity improves it (Xu et al., 2023). Fewer spines on these neurons could therefore weaken the prefrontal “brake” on emotional responses, which would fit with the weaker functional connectivity between the prefrontal cortex and the amygdala reported in people who experienced childhood maltreatment (Teicher et al., 2016). Importantly, in adult rodents the effects of stress, even chronic stress, generally reverse within a few weeks once the stress ends (Lupien et al., 2009). In the rat mPFC, for example, apical dendrites shortened by three weeks of daily restraint stress recover after three weeks without stress (Radley et al., 2005). By contrast, the loss we observed persisted long after ELS had ended. One possible reason is that stress hormones act on spines that are still being established: in mice, chronic glucocorticoid excess leads to the loss of stable spines formed early in life (Liston and Gan, 2011), so stress during this window may remove spines that would otherwise have been kept into adulthood. This suggests that early adversity may leave a more lasting structural mark on the prefrontal cortex than comparable stress later in life. Because we did not measure behavior, these functional interpretations remain hypotheses.

### 4.5 Considerations and future directions

This study was intended as an initial survey of spine density across the PL, IL and BLA after ELS, with the aim of identifying effects to test in a larger sample. Several features support the main finding: all three regions were sampled in the same animals, spines were counted blind to group, the animal rather than the neuron or segment was the unit of analysis, and the effect of ELS was large (η^2^G = .60) and in the same direction in all three regions. At the same time, the sample was small, with three male mice per group drawn from two cohorts and two litters per group. A sample of this size can detect only large effects and cannot separate litter effects from the effect of ELS (Lazic and Essioux, 2013). The regional and dendritic patterns we describe were not supported by significant interactions and rest on exploratory, uncorrected comparisons, so they should be treated as hypotheses rather than conclusions. Our sampling and measures were also focused. We examined proximal oblique branches only, and because stress-related spine loss in the mPFC can be greatest on distal apical segments (Radley et al., 2006; Xu et al., 2023), the full extent of apical spine loss after ELS may be larger than we measured. We measured spine density alone, without spine shape, dendritic branching or behavior, and because ELS effects on the prefrontal cortex can differ in females (Farrell et al., 2016), the findings may not extend to them. Without measures of dendritic length, we also cannot tell whether fewer spines per 10 µm means fewer spines overall; if ELS also lengthens dendrites, as reported for IL apical dendrites in adolescent females (Farrell et al., 2016), density could fall while the total number of spines changes less. Absolute spine densities also differ widely across studies because of differences in labeling, imaging and the dendritic segments sampled; for example, Xu et al. (2023) averaged spine density across the whole dendritic tree, including spine-sparse segments, so our values are best compared between groups within this study rather than with values reported elsewhere.

These open questions define the next steps. The effect sizes reported here provide a basis for powering a larger study that includes both sexes and more litters, samples distal as well as proximal apical dendrites, examines spine types, records maternal care and pup weight during the ELS window, and relates these structural changes to prefrontal-dependent behaviors such as fear extinction.

## 6 ACKNOWLEDGMENTS

We thank Dr. Ivan Hernandez for training in microscopy, and the members of the laboratory, Adam Pittner, Esther Gonzalez and Kelsie Childress, for their assistance with the early life stress procedures and Golgi-Cox staining. We also thank the staff of the Division of Comparative Medicine at SUNY Downstate Health Sciences University for animal care.

